# Brassinosteroids Influence Arabidopsis Hypocotyl Graviresponses Through Changes In Mannans And Cellulose

**DOI:** 10.1101/557777

**Authors:** Marc Somssich, Filip Vandenbussche, Alexander Ivakov, Norma Funke, Colin Ruprecht, Kris Vissenberg, Dominique Van Der Straeten, Staffan Persson, Dmitry Suslov

## Abstract

The force of gravity is a constant environmental factor. Plant shoots respond to gravity through negative gravitropism and gravity resistance. These responses are essential for plants to direct the growth of aerial organs away from the soil surface after germination and to keep an upright posture above ground. We took advantage of the effect of brassinosteroids on the two types of graviresponses in *Arabidopsis thaliana* hypocotyls to disentangle functions of cell wall polymers during etiolated shoot growth. The ability of etiolated Arabidopsis seedlings to grow upwards was suppressed in the presence of 24-epibrassinolide (EBL) but enhanced in the presence of brassinazole (BRZ), an inhibitor of brassinosteroid biosynthesis. These effects were accompanied by changes in cell wall mechanics and composition. Cell wall biochemical analyses and confocal microscopy of the cellulose-specific pontamine S4B dye revealed that the EBL and BRZ treatments correlated with changes in cellulose fibre organization and mannan content. Indeed, a longitudinal re-orientation of cellulose fibres supported upright growth whereas the presence of mannans reduced gravitropic bending. The negative effect of mannans on gravitropism is a new function for this class of hemicelluloses, highlighting evolutionary adaptations by which aquatic ancestors of terrestrial plants colonized land.

## Introduction

All growing plant cells are encased in primary cell walls, which are strong to maintain cell and tissue integrity, yet extensible to allow for growth (Cosgrove 2016). How these conflicting properties are achieved is uncertain because the exact architecture of primary cell walls is not well defined (Cosgrove 2018). The traditional cell wall model, in which a load-bearing network of cellulose microfibrils, cross-linked by hemicelluloses, is embedded into an amorphous matrix of pectins and structural glycoproteins (Carpita and Gibeaut 1993), has been substantially modified and improved (Cavalier et al. 2008; Dick-Pérez et al. 2011). These modified models include a wall load-bearing capacity that is defined by either lateral cellulose/xyloglucan/cellulose contacts in restricted areas called “biomechanical hotspots” (Park and Cosgrove 2012), and/or cellulose/pectin interactions (Phyo et al. 2017a; Phyo et al. 2017b), and/or arabinogalactan proteins covalently linked with pectin and arabinoxylans (Tan et al. 2013). Additionally, cellulose microfibrils are packed into macrofibrils in primary cell walls (Anderson et al. 2010), the effect of which on the wall properties is yet to be determined. At present, there is little consensus on the role of particular polysaccharides in cell walls, and there is therefore a need for experimental models that could help understand their functions and interactions.

Plant shoots normally grow upright, against the gravity vector, which is supported by two principal graviresponses – gravitropism and gravity resistance. Gravitropism is defined as directed growth of a plant or plant organ in response to gravity (Kiss 2000). Negative shoot gravitropism takes a form of upward bending, an asymmetric shoot growth that restores its vertical position after inclination (Morita 2010). This response requires flexibility of cell wall polymers and is affected by their compression resistance on the concave side of the gravitropically bending organ. As such, gravitropism is a useful model for studying these rarely addressed properties of wall polymers. Shoot gravity resistance is defined as mechanical resistance to the gravitational force (Hoson and Wakabayashi 2015). During upright shoot growth, supported by gravity resistance, the conflicting properties of growing walls are especially prominent: their extensibility should be delicately balanced with the strength not only to maintain cell and tissue integrity, but also to carry the organ’s weight in the field of gravity. The fine balance of the opposite wall properties in this system implies that even minor experimentally imposed modifications of cell wall polymers will be translated into changes of shoot posture. Hence, the upright shoot growth may work as a sensitive model for establishing the structural basis of conflicting wall properties.

Brassinosteroids (BRs) constitute a class of phytohormones (Singh and Savaldi-Goldstein 2015) that impact graviresponses of Arabidopsis hypocotyls that are largely built of primary cell walls (Vandenbussche et al. 2011). The gravity resistance of hypocotyls was suppressed by epibrassinolide (EBL), one of the most active natural BRs, while brassinazole (BRZ), a specific inhibitor of BR biosynthesis, stimulated their gravitropism. These effects did not result from modified gravity perception but were accompanied by changes in cell wall mechanics (Vandenbussche et al. 2011). The biochemical basis of the BR-induced alterations in wall physical properties is, however, largely unknown. Comprehensive microarray data on BR effects on Arabidopsis plants revealed changes in many dozens of cell wall-related genes (Goda et al. 2002; Song et al. 2009; Sun et al. 2010; Yu et al. 2011). Thus, the action of BRs, including that of EBL and BRZ, on Arabidopsis hypocotyl graviresponses could be mediated by a number of modifications at the cell wall level.

In the present work we used the strong EBL and BRZ effects on Arabidopsis hypocotyl graviresponses to better understand cell wall polysaccharide functions employing high-resolution confocal microscopy, biochemical and biomechanical analyses.

## Results

### Brassinosteroids strongly affect etiolated shoot graviresponses in Arabidopsis

To corroborate that the graviresponses of etiolated Arabidopsis seedlings were altered upon changes in BR signaling (Vandenbussche et al. 2011), we grew seedlings on horizontal Petri plates in the presence of exogenous EBL or BRZ and counted upright hypocotyls (Fig. 1). Plants grown on one-half strength MS media had 40-80% of upright hypocotyls (Fig. 1A). Media supplemented with EBL (100 nM) resulted in a decreased proportion of standing hypocotyls to 0-5% (Fig. 1B), while media containing BRZ (1 μM) increased it to 90-100% (Fig. 1C). These observations were consistent essentially from the moment of germination and demonstrated that BRs suppressed normal shoot graviresponses.

**Fig. 1.**
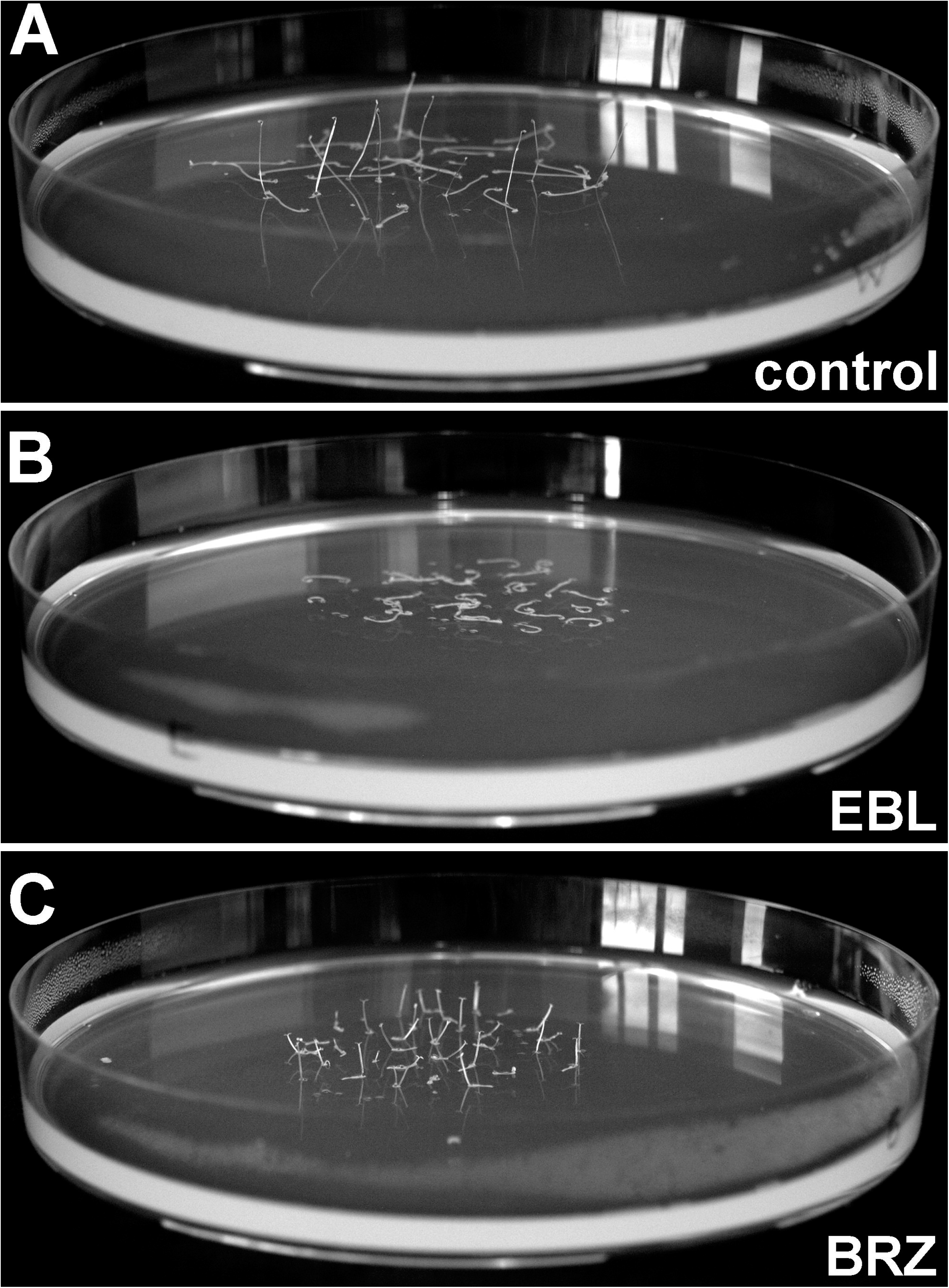
Brassinosteroids affect upright growth of hypocotyls in Arabidopsis plants. Etiolated *Col-0* seedlings were grown on horizontal Petri plates with one-half strength agarized Murashige and Skoog medium without supplements (A), with 100 nM EBL (B) or 1 μM BRZ (C). Five-day-old plants were photographed.

### Brassinosteroids influence etiolated shoot graviresponses via changes in cell wall mechanics

The influence of EBL on the hypocotyl posture (Fig. 1B) was hypothesized to emanate from cell wall weakening that resulted in a loss of hypocotyl gravity resistance and their inability to stand upright in the field of gravity (Vandenbussche et al. 2011). EBL had no effect on gravitropic bending in Arabidopsis plants grown on vertical Petri plates and gravistimulated by a 90 degrees clockwise rotation of the plates (see Fig. 4 in Vandenbussche et al. 2011). In contrast, BRZ increased both the percentage of upright hypocotyls and their gravitropism, but its effect on cell wall mechanics remained unclear (Vandenbussche et al. 2011).

**Fig. 2.**
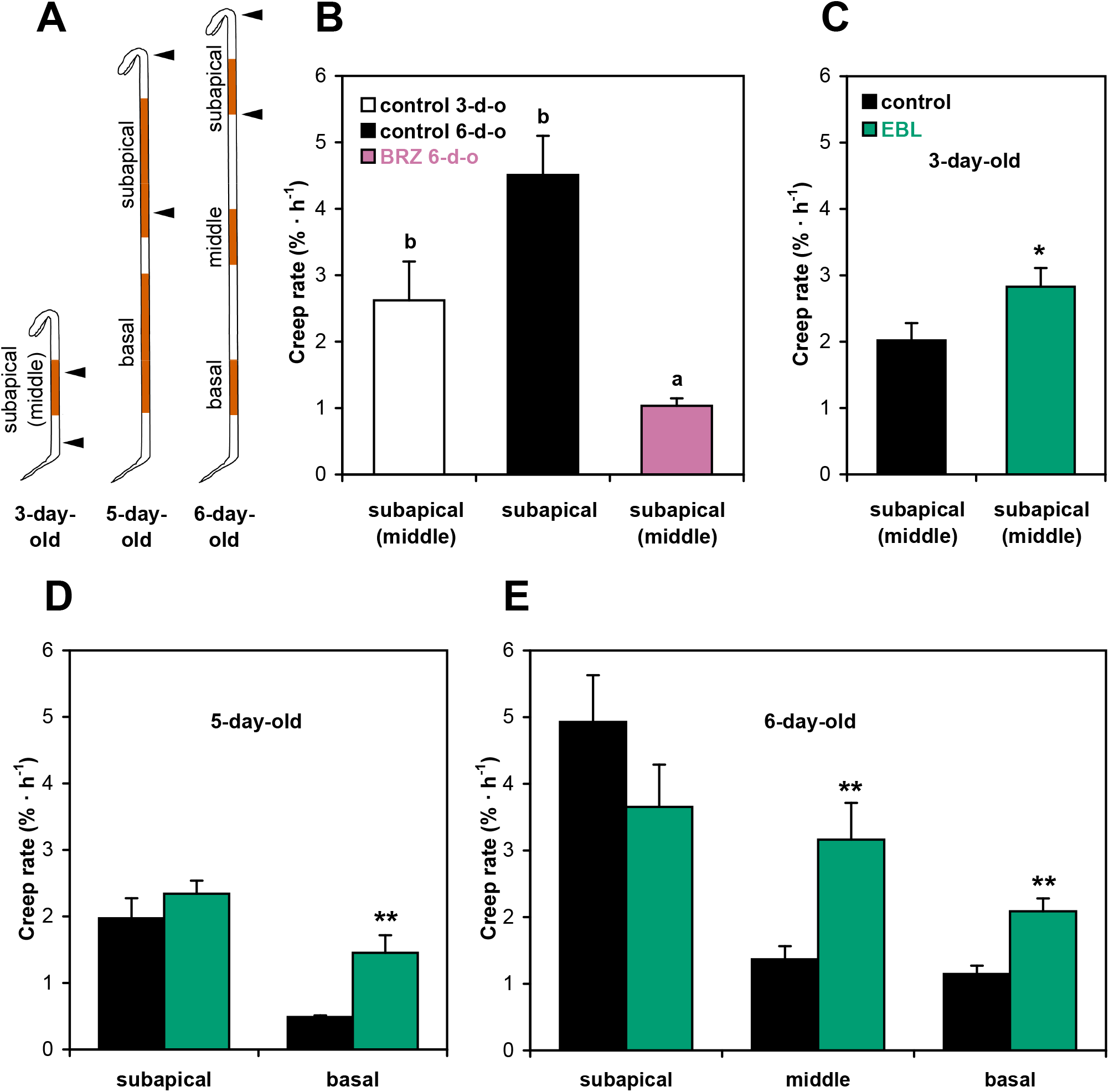
Brassinosteroids change cell wall mechanics in Arabidopsis hypocotyls. (A) The schematic representation of regions extended by the creep method (orange shadings) in hypocotyls of different age. Arrowheads mark borders of growing zones in hypocotyls of *Col-0* plants of the respective age. Changes in creep rates by BRZ (1 μM) in hypocotyls of 6-day-old *Col-0* plants (B) and by EBL (100 nM) in hypocotyls of 3-day-old (C), 5-day-old (D) and 6-day-old (E) *Col-0* plants. All seedlings in B-E were grown on horizontal Petri plates. Creep rates were measured under 750 mg (B, C, E) or 625 mg (D) loads. Data are means ±SE (n=10). Different letters in (B) mark significant differences (*P* < 0.05) revealed by Games-Howell’s post-hoc test performed after ANOVA. Asterisks in (C-E) denote significant difference (\**P* < 0.05; \*\**P* < 0.01) between the respective zones of EBL-grown and control seedlings determined by Student’s *t*-test.

**Fig. 3.**
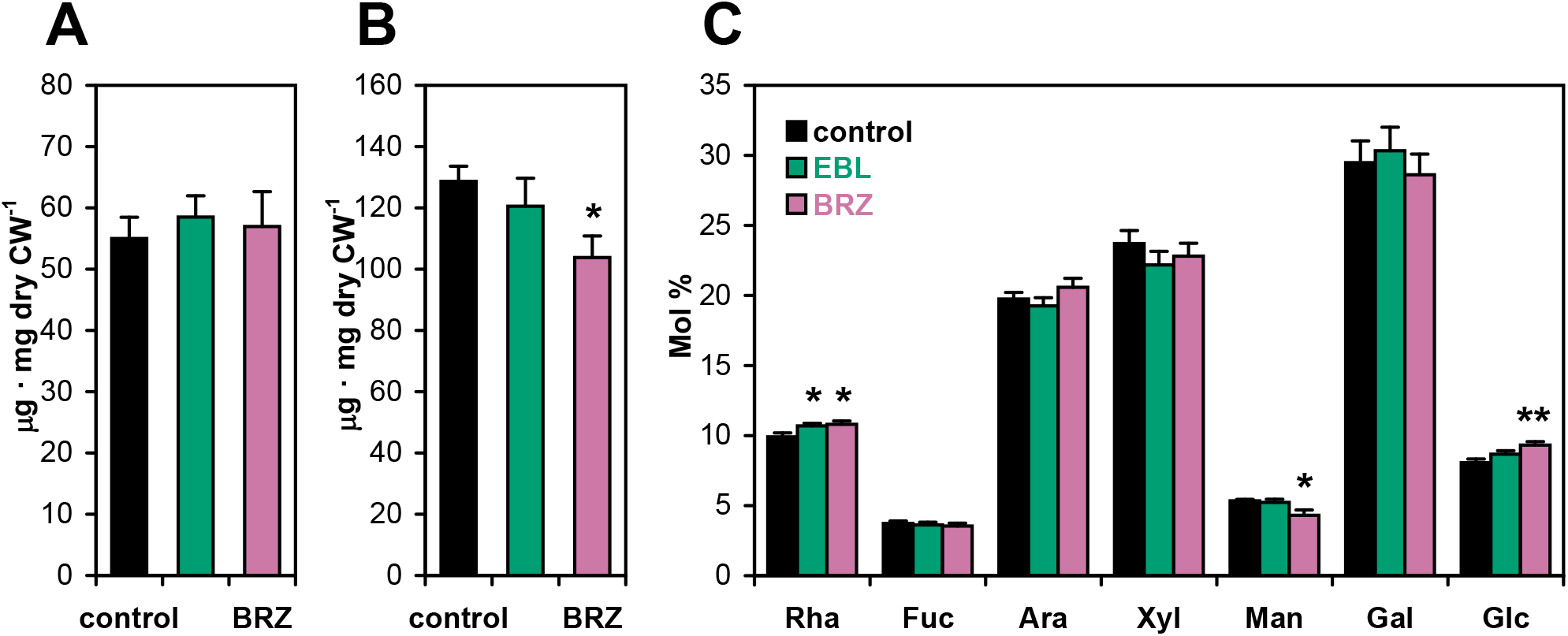
Biochemical composition of cell walls in Arabidopsis plants as affected by brassinosteroids. The levels of uronic acids (A), crystalline cellulose (B) and the monosaccharide composition of cell wall matrix polymers (C) were determined in 5-day-old etiolated *Col-0* seedlings grown on horizontal Petri plates without supplements in the growth medium (control), in the presence of 1 μM BRZ or 100 nM EBL. Data are means ±SE (n=9). Asterisks denote significant differences (\**P* < 0.05; \*\**P* < 0.01) revealed by Dunnett’s post-hoc test performed after ANOVA.

**Fig. 4.**
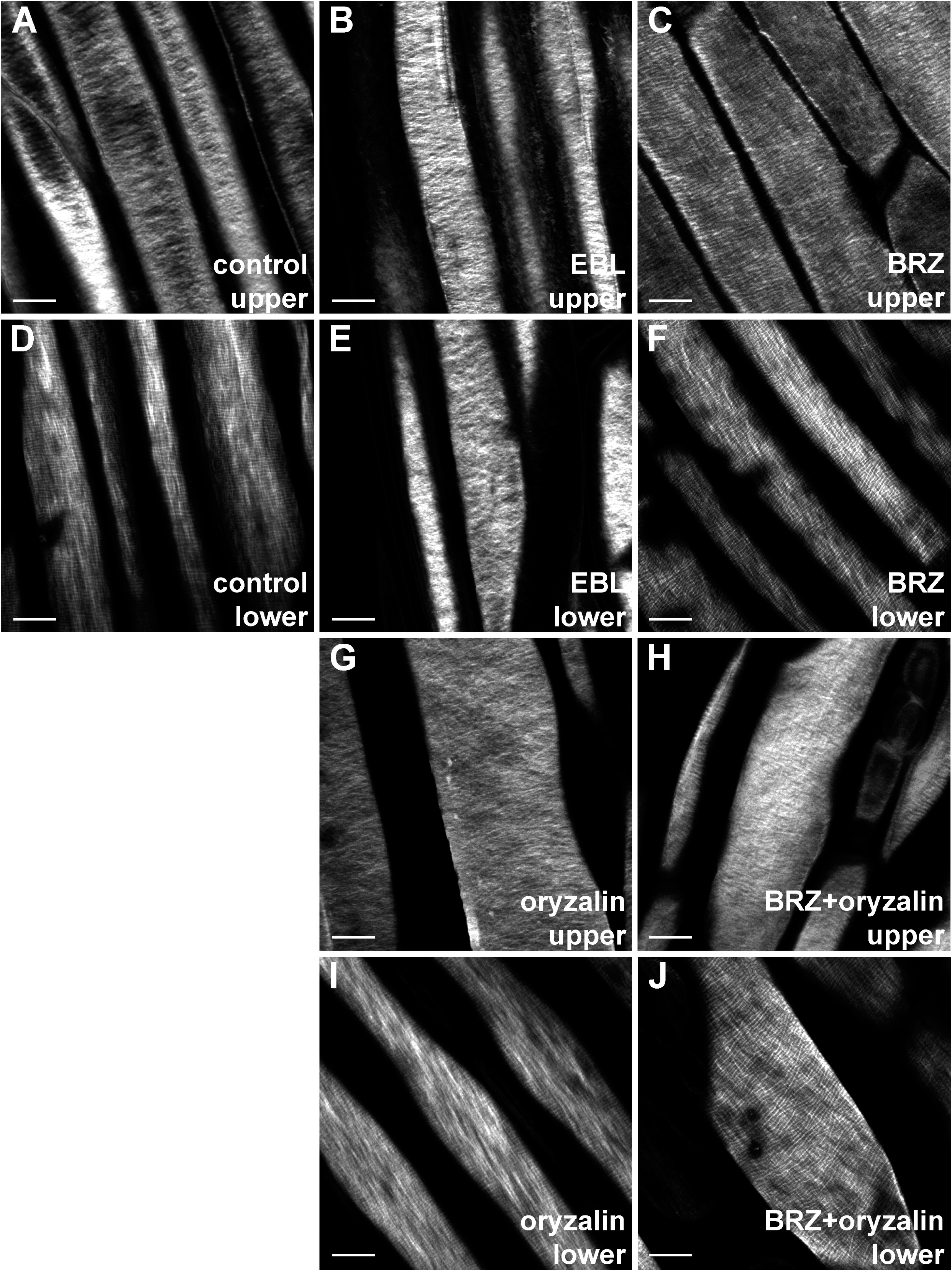
Brassinosteroid and oryzalin effects on the arrangement of cellulose macrofibrils in the outer epidermal cell wall of hypocotyls from 5-day-old etiolated *Col-0* Arabidopsis plants grown on horizontal Petri plates. Hypocotyls were extracted under mild conditions, their walls were stained with Pontamine Fast Scarlet 4B, the cellulose-specific fluorescent dye, and observed under a super-resolution Airyscan confocal microscope. Cellulose orientation in projections of three-five optical sections across the whole wall thickness is shown (A) in the upper growing and (D) the lower non-growing region of hypocotyls from control seedlings grown on one-half strength agarized Murashige and Skoog medium without supplements; (B) the upper and (E) the lower region of hypocotyls from plants grown in the presence of 100 nM EBL; (C) the upper and (F) the lower region of hypocotyls from plants grown in the presence of 1 μM BRZ; (G) the upper and (I) the lower region of hypocotyls from plants grown in the presence of 250 nM oryzalin; (H) the upper and (J) the lower region of hypocotyls from plants grown in the presence of 1 μM BRZ and 250 nM oryzalin. Scale bars are 10 μm.

We used the creep (constant-load) method to assess the shoot gravity resistance. Whenever hypocotyl length afforded, we stretched their basal regions, carrying the main part of the organ weight in the field of gravity, as well as their apical growing regions that are responsible for gravitropic bending (Fig. 2A). The scheme in Fig. 2A illustrates the established age-dependent shift of growth zones in etiolated Arabidopsis hypocotyls; from the base towards cotyledons (Gendreau et al. 1997; Bastien et al. 2016). As BRZ greatly inhibited etiolated hypocotyl elongation (Fig. 1A, C), we compared cell wall creep in BRZ-grown plants with their age-matched 6-day-old untreated counterparts, as well as with 3-day-old untreated “genetic controls”, which are approximately the same length as the 6-day-old BRZ-grown plants (Fig. 2B). BRZ significantly decreased creep rates compared with both controls, which is consistent with cell wall strengthening. Thus, the inhibition of BR biosynthesis may increase the percentage of upright hypocotyls, not only through the stimulation of gravitropic bending (Fig. 4 in Vandenbussche et al. 2011), but also by increasing their gravity resistance (Fig. 2B). The creep rate analyses in EBL-grown plants revealed cell wall weakening that was mostly restricted to the basal nonexpanding hypocotyl zones (Fig. 2C-E). Hence, EBL may affect the posture of etiolated hypocotyls (Fig. 1B) by decreasing their gravity resistance in the basal regions (Fig. 2D-E) where cell expansion has already ceased (Gendreau et al. 1997).

### Changes in cell wall biochemistry underpin the effects of BRs on graviresponses

To investigate how the cell walls were altered during BR exposure, thereby influencing cell wall mechanics and the hypocotyl posture, we performed standard cell wall biochemical analyses (Fig. 3). The content of uronic acids, which are the principal constituents of pectins, was unaffected by the EBL or BRZ treatments (Fig. 3A). By contrast, crystalline cellulose levels were significantly decreased in BRZ treated seedlings as compared with the untreated control, while EBL had no effect on this polymer level (Fig. 3B). As for monosaccharide composition of cell wall matrix polymers, both EBL and BRZ treatments significantly increased rhamnose content (Fig. 3C). In addition, BRZ treatment decreased mannose and increased glucose in cell wall matrix polysaccharides (Fig. 3C). Changes in rhamnose are likely due to the metabolism of rhamnogalacturonan I, which is the main source of this monosaccharide in primary cell walls. As EBL and BRZ caused opposite effects on the hypocotyl posture (Fig. 1B, C) but induced very similar increases in rhamnose (Fig. 3C), we argued that it is unlikely that this sugar is underpinning our observed phenotype. We did therefore not consider the rhamnose change in further details. Mannose is the principal constituent of cell wall mannans and the BRZ-induced decrease of mannose could therefore be caused by partial mannan depletion in the seedling cell walls. The BRZ-induced increase in glucose was not accompanied by any xylose accumulation (Fig. 3C) indicating that it was not associated with xyloglucans. On the other hand, the increase in glucose correlated with a decrease in crystalline cellulose (Fig. 3B). The inverse relationship between crystalline cellulose (Fig. 3B) and glucose (Fig. 3C) could be explained by a larger amorphous component of cellulose microfibrils, which is likely to be sensitive to TFA hydrolysis, perhaps explaining the apparent increase in glucose content in BRZ treated seedlings (Fig. 3C).

### EBL effect on the posture of etiolated hypocotyls is related to cellulose arrangement

Rather unexpectedly, the strong effect of EBL on cell wall mechanics (Fig. 2C-E) was not associated with prominent changes in cell wall biochemical composition (Fig. 3). Nevertheless, not only the levels of certain cell wall polymers, but also their orientations and interactions in the wall affect cell wall mechanics, with implications both on cellular strength and expansion. This is especially true for cellulose, the strongest cell wall component (Suslov and Verbelen 2006; Suslov et al. 2009). To examine the orientation of cellulose in outer epidermal cell walls of hypocotyls from control and EBL-grown plants we used the specific fluorescent dye Pontamine Fast Scarlet 4B (S4B) (Anderson et al. 2010). We imaged the dye-associated cellulose fibers using spinning-disc or high-resolution Airyscan confocal microscopy. With these setups we could reveal distinct cellulose macrofibril orientations. To improve the penetration of the dye and visualization of the cellulose fibers along the whole hypocotyl length we performed a mild extraction of cell walls. Without this treatment the cellulose fibers were not well discerned in the basal parts of living hypocotyls. Nevertheless, living and extracted seedlings displayed similar cellulose arrangements in the upper growing region of their hypocotyls. In this part of control seedlings transverse buckling was frequently observed in the innermost wall layer. This phenomenon occurred in approximately 50% of both living (Supplementary Fig. S1) and extracted control hypocotyls, and complicated cellulose macrofibril visualization on the inner wall face. In the remaining samples, i.e. in which we did not observe the buckling, cellulose macrofibrils were clearly transverse in the innermost wall layer and longitudinal in the outermost layer (Supplementary Video S1; Fig. 4A; Table 1). In the basal non-growing region of control hypocotyls, we observed slight or no buckling of the wall inner surface. These walls contained less transverse macrofibrils in the innermost layer and thicker longitudinal macrofibrils in the outermost layer compared with the upper growing region of hypocotyls (Supplementary Video S2; Fig. 4D; Table 1). In the upper region of approximately 50% of hypocotyls from EBL-grown plants, we observed irregular buckling in the innermost wall. In the samples without buckling, cellulose macrofibrils were more obliquely oriented (Fig. 4B), or decreased in relative abundance, compared with the untreated control. EBL did not influence longitudinal macrofibrils in the outermost wall layer in the growing zone of hypocotyls (Supplementary Video S3). The most dramatic EBL effect on cellulose orientation was found in the basal region of hypocotyls where it essentially eliminated longitudinal macrofibrils, such that the remaining ones were apparently thinner and had either oblique or random orientation (Fig. 4E; Supplementary Video S4; Table 1). This reduction in longitudinal macrofibrils at the hypocotyl base was a unique effect observed only in EBL-treated seedlings (Table 1; Fig. 4).

**Table 1.** Overview of cellulose macrofibril orientations Cellulose macrofibril orientation across the whole thickness of outer epidermal cell walls of Arabidopsis hypocotyls was determined using S4B dye, spinning-disc and super-resolution confocal microscopy. The prevalence of four types of cellulose macrofibril orientation across the wall thickness was characterized by scores 1 to 3 as high (3), intermediate (2) or low (1). These values were averaged between all cells analyzed for a given treatment and presented as means ± SD.

| Treatment | Cellulose orientation |  |  |  | Number of repeats<sup>*</sup> |
| --- | --- | --- | --- | --- | --- |
|  | transverse | oblique | random | longitudinal |  |
| Col-0, upper<sup>†</sup> | 1.82±0.70 | 0.09±0.28 | 0.08±0.25 | 1.43±0.63 | 38 |
| Col-0, lower<sup>‡</sup> | 1.18±0.56 | 0.16±0.37 | 0.05±0.27 | 1.89±0.51 | 38 |
| Col-0+EBL, upper | 0.42±0.69 | 0.47±0.77 | 0.10±0.30 | 1.53±0.61 | 19 |
| Col-0+EBL, lower | 0.33±0.55 | 1.40±0.72 | 0.43±0.63 | 0.63±0.56 | 30 |
| Col-0+BRZ, upper | 1.48±0.60 | 0.02±0.09 | 0.26±0.44 | 1.82±0.67 | 31 |
| Col-0+BRZ, lower | 0.65±0.49 | 0.09±0.29 | 0.17±0.39 | 1.91±0.29 | 23 |
| Col-0+ory, upper | 0.90±0.84 | 0.52±0.70 | 1.22±0.85 | 1.56±0.73 | 25 |
| Col-0+ory, lower | 0.23±0.44 | 0.15±0.38 | 0.77±0.60 | 2.00±0.00 | 13 |
| Col-0+ory+BRZ, upper | 0.63±0.66 | 0.22±0.36 | 1.13±0.67 | 1.32±0.75 | 26 |
| Col-0+ory+BRZ, lower | 0.20±0.41 | 0.11±0.32 | 1.20±0.70 | 1.90±0.31 | 20 |

\*: Each repeat is an outer cell wall from an individual epidermal cell. The data are based on three-four independent experiments with at least four seedlings analyzed in each of them. <sup>†</sup>: upper region of a hypocotyl. <sup>‡</sup>: lower region of a hypocotyl. EBL: epibrassinolide; BRZ: brassinazole; ory: oryzalin.

To confirm that the EBL effect on hypocotyl posture was associated with reduced macrofibril organization, we attempted to “randomize” cellulose orientation in the cell walls and then see how this affected the percentage of standing hypocotyls (Fig. 5). For this purpose, Arabidopsis seedlings were grown in the presence of 250 nM oryzalin, which partially disassembles cortical microtubules, thereby affecting the direction of cellulose microfibril deposition in the cell wall (Paredez et al. 2006). This low concentration of oryzalin induced similar changes in cellulose arrangement as those observed with EBL in the upper part of hypocotyls (compare Supplementary Video S5 with Supplementary Video S3; and Fig. 4B with 4G; Table 1), and significantly decreased the percentage of standing hypocotyls (Fig. 5). Thus, intact cellulose orientation is important for keeping the hypocotyls upright, and the mechanism of EBL action on their posture may be based on changes in cellulose arrangement. However, the effect of oryzalin on the percentage of upright hypocotyls (Fig. 5) was weaker than that of EBL (Fig. 1B). This difference can be explained by the inability of oryzalin to alter longitudinal cellulose macrofibrils in the basal region of hypocotyls, in contrast to the effect of EBL (compare Supplementary Video S6 with Supplementary Video S4; and Fig. 4E with Fig. 4I; Table 1). These longitudinal macrofibrils accumulated in the outermost wall layer where they could contribute to the mechanical strength and upright growth of hypocotyls. Their reduced presence in EBL-treated seedlings (Fig. 4E, Supplementary Video S4) correlated with weaker walls at the base of hypocotyls (Fig. 2D, E).

**Fig. 5.**
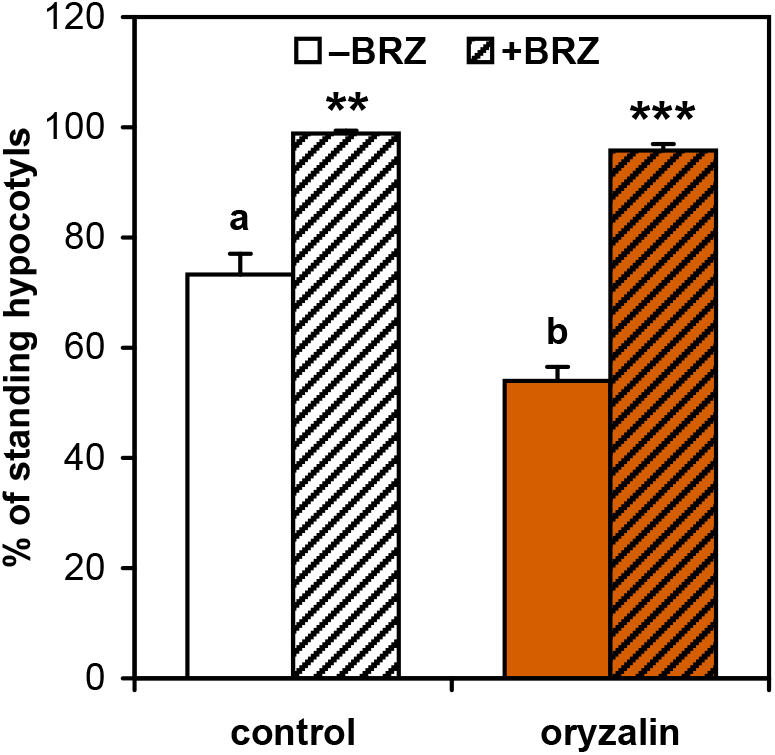
Cellulose orientation affects the percentage of standing Arabidopsis hypocotyls. *Col-0* seedlings were grown in darkness on horizontal Petri plates for 5 days without supplements (white bar), with 250 nM oryzalin (orange bar), in each case with 1 μM BRZ (hatched bars) or without (solid color bars). Data are means ±SE (n=6). Different letters denote the significant difference between control and oryzalin-grown plants (*P*=0.0017; Student’s *t*-test). Asterisks indicate significant effects of BRZ in control or oryzalin-grown seedlings (\*\**P* < 0.01; \*\*\**P* < 0.001; Student’s *t*-test).

### BRZ increases the percentage of upright etiolated hypocotyls via several different mechanisms

Interestingly, BRZ increased the percentage of upright hypocotyls independently of oryzalin treatment (250 nM) (Fig. 5). These data indicate that BRZ either regulates graviresponses somewhat independently of cellulose orientation, and/or antagonizes the oryzalin-induced macrofibril randomization. Our microscopic observations supported the first option, because we observed less ordered cellulose orientations in the upper parts of seedlings treated with oryzalin and BRZ (Fig. 4C, H; Supplementary Videos S7, S8; Table 1). At the same time the cellulose arrangement in the basal hypocotyl part was essentially similar in the untreated controls and seedlings grown in the presence of BRZ, oryzalin, and BRZ plus oryzalin (Fig. 4D, F, I, J; Supplementary Videos S2, S6, S9, S10; Table 1).

Thus, our imaging data indicate that BRZ may affect graviresponses via mechanisms not related to cellulose. One such mechanism might be mediated by reducing the mannan content (Fig. 3C). To assess whether a decrease in mannan content affected the graviresponses we studied triple *csla2csla3csla9* mutant plants lacking detectable glucomannans in mature Arabidopsis stems (Goubet et al. 2009). Transcriptomics data indicate that out of the nine *CSLA* genes in Arabidopsis, *CSLA2*, *3*, and *9* have the highest expression in hypocotyls (Winter et al. 2007). The *csla2csla3csla9* triple mutation did not influence the percentage of standing hypocotyls (Supplementary Fig. S2). However, by analogy with BRZ, the mutants displayed significantly accelerated gravitropic bending in reorientation assays with plants grown on vertical Petri plates (Fig. 6; Supplementary Fig. S3). Interestingly, the coefficient of BRZ stimulation of gravitropic bending was reduced in the background of *csla2csla3csla9* compared with Col-0 (Fig. 6, Table 2). The above findings show that the decreased mannan content (Fig. 3C) accounts – at least in part - for the BRZ-induced increase in gravitropic bending.

**Fig. 6.**
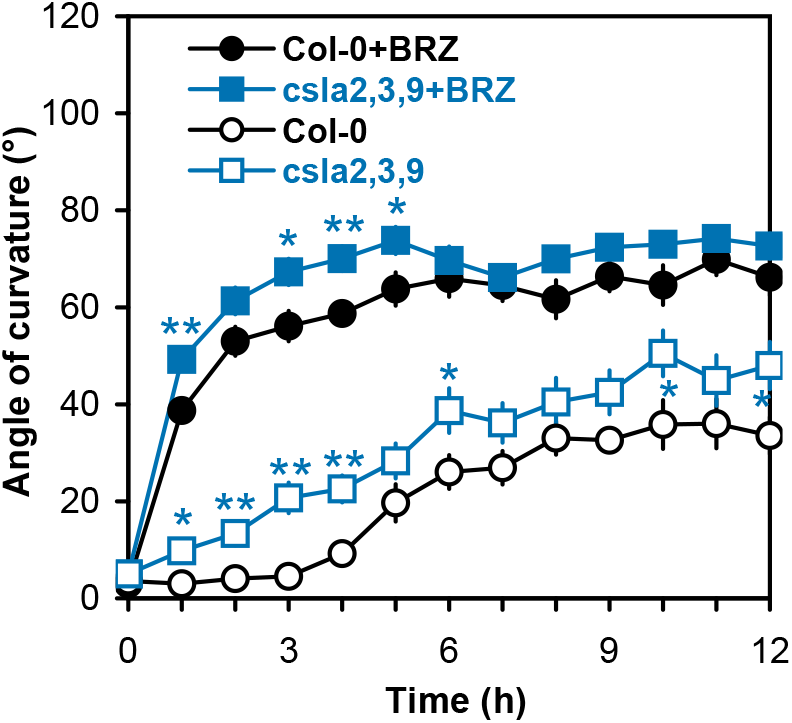
Hypocotyls of glucomannan-deficient *csla2csla3csla9* mutant plants demonstrate increased gravitropic bending. Etiolated *Col-0* and *csla2csla3csla9* mutant seedlings were grown on vertical Petri plates with or without 1 μM BRZ for 3 days, subsequently gravistimulated by a 90-degree clockwise rotation of the plates, and gravitropic bending of their hypocotyls was followed by an infrared imaging system for 12 h. Data are means ±SE (n=10). Asterisks denote significant differences (\**P* < 0.05; \*\**P* < 0.01; Student’s *t*-test) between *csla2csla3csla9* and *Col-0* plants.

**Table 2.**
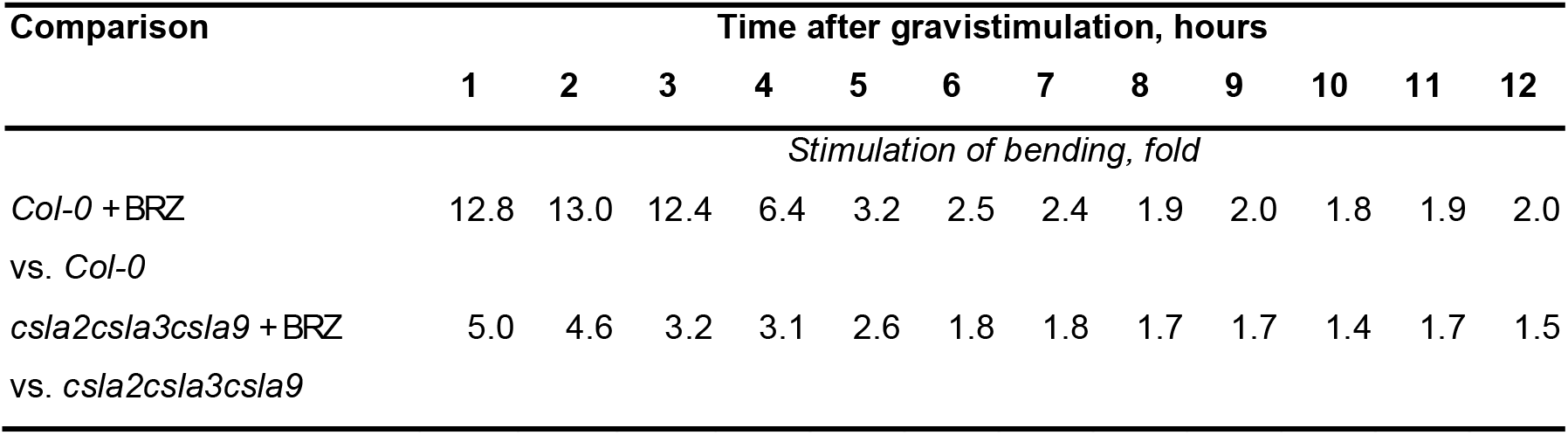
Stimulation of gravitropic bending by brassinazole is impaired in the triple *csla2csla3csla9* mutant compared with *Col-0* plants Gravitropic bending in the presence of BRZ (Fig. 6) was divided by that without BRZ for *Col-0* and *csla2csla3csla9*, respectively.

Plant cell expansion is known to be induced by cell wall loosening (Cosgrove 2016; Ivakov et al. 2017). Cell growth is slow and uniform along the length of young etiolated hypocotyls that have just emerged from germinating seeds (Refrégier et al. 2004). A wave of rapid cell expansion starts at the base of hypocotyls at about 48 h post induction, after which it spreads acropetally towards cotyledons (Gendreau et al. 1997; Bastien et al. 2016). Hence, cell wall loosening is highly induced in the basal region of hypocotyls carrying their main weight, which can interfere with keeping the upright position of this organ in the field of gravity. As the effect of BRZ on hypocotyl posture is visible from the moment of emergence from the seed, it may be caused by changes in cell expansion and, hence, cell wall loosening in very young hypocotyls. To test this option, cell length was measured in epidermal cell files at two developmental stages (Fig. 7A), allowing growth rate calculation for individual cells along hypocotyls (Fig. 7B). Only basal halves of epidermal cell files along hypocotyls (ten lower cells) were considered. These are the regions where the rapid growth is first initiated in hypocotyls (Gendreau et al. 1997). Cell length was measured at 55 h post induction, the time point corresponding to the earliest phase of rapid growth in hypocotyls (Pelletier et al. 2010), and at 72 h post induction, when maximal growth rate is attained (Gendreau et al. 1997). Epidermal cell length was significantly lower in hypocotyls of BRZ-treated vs. control plants for all cells with the exception of the first cell at 55 h illustrating severe growth inhibition due to a decrease in BR biosynthesis (Fig. 7A). The rate of cell expansion was also significantly lower for the majority of cells in BRZ-treated vs. control plants indicating that the wall loosening was inhibited in the former (Fig. 7B). Interestingly, two peaks of growth inhibition were observed in BRZ-treated seedlings: at the base of hypocotyls (cells one and three) and in their most apical zone examined (cells nine and ten) (Fig. 7B). Thus, the wave of rapid growth is initiated in a higher region of hypocotyls and spreads acropetally slower in BRZ-treated vs. control plants. Overall, the above data indicate that cell wall loosening is inhibited more strongly at the base of hypocotyls, i.e. in the region responsible for carrying the seedling weight in the field of gravity. This inhibition can be another mechanism by which BRZ increases the percentage of standing hypocotyls.

**Fig. 7.**
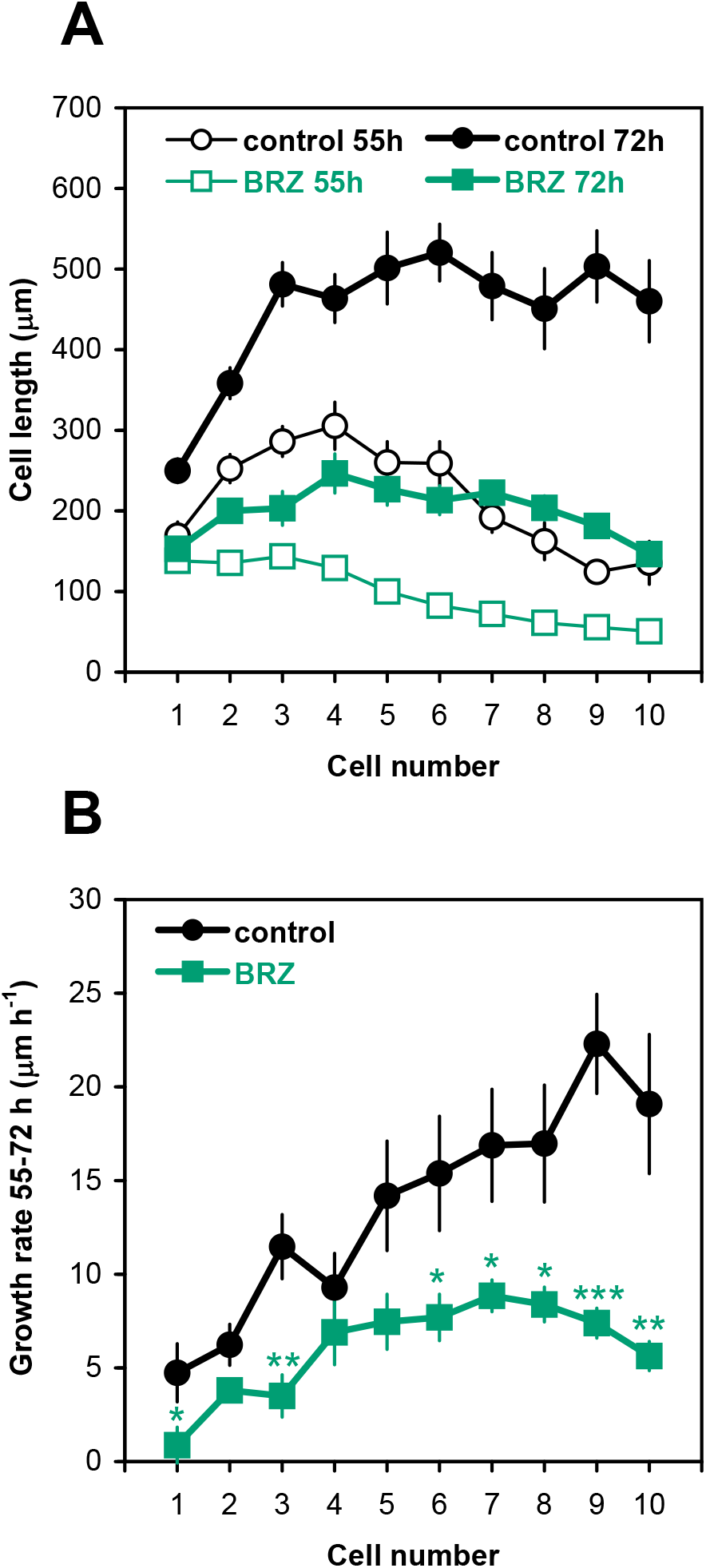
BRZ inhibits cell expansive growth at the base of Arabidopsis hypocotyls. (A) Epidermal cell length distribution along the lower parts of hypocotyls in 55- and 72-hour-old etiolated *Col-0* seedlings grown on horizontal Petri plates with 1 μM BRZ or without (control). Cells are numbered from the base of hypocotyls towards cotyledons. Data are means ±SE (n=9). All the cells are significantly shorter in the presence of BRZ than the respective control cells (*P* < 0.01; Student’s *t*-test) with the exception of cells 1 in 55-hour-old plants (not shown on the plot A). (B) Average cell expansion rates at the base of hypocotyls calculated from the data of plot A (means ±SE, n=9). Asterisks mark significant differences (\**P* < 0.05; \*\**P* < 0.01; \*\*\**P* < 0.001; Student’s *t*-test).

## Discussion

The two prerequisites for upright growth of young shoots, mechanical strength and gravitropic bending, are based on distinct physiological mechanisms, which are linked through the cell wall characteristics. The mechanical strength of young shoots is defined by turgor (Shah et al. 2017). This is well illustrated by wilting, a loss of turgor, upon which young shoots fall down on the ground. Turgor depends on the transmembrane concentration gradient of osmotically active compounds, the hydraulic conductance and wall yielding properties (Ray et al. 1972). The equation for turgor (Ray et al. 1972) implies that cell wall weakening (Fig. 2C-E) and strengthening (Fig. 2B) in the presence of EBL and BRZ, respectively, will contribute to lower and higher turgor values, which could underpin the BR effects on hypocotyl posture (Fig. 1). However, we cannot exclude that these effects are partially mediated by osmotic adjustments also influencing turgor pressure, which was not addressed here. Similarly, the selective BRZ-induced growth inhibition at the hypocotyl base (Fig. 7B), presumably mediated by decreased cell wall yielding, could elevate turgor, thereby maintaining their upright position.

Gravitropic bending results from asymmetric cell expansion on the opposite flanks of gravistimulated organs (Miller et al. 2007). In young shoots put horizontally this response originates from simultaneous growth inhibition and stimulation on their upper and lower flanks, respectively (Bagshaw and Cleland 1990; Cosgrove 1990a; Ikushima et al. 2008). As in the case of cell expansion in vertical plant organs (Cosgrove 2018), the growth responses during gravitropic bending are controlled by cell wall yielding properties (Bagshaw and Cleland, 1990; Cosgrove 1990b; Edelmann and Samajova 1999; Ikushima et al. 2008). Cell wall extensibility always decreases on the upper sides and sometimes increases on the lower sides of gravistimulated shoots, such that the overall cell wall loosening or tightening prevails depending on the species (Edelmann 1997) and/or the phase of gravitropic bending (Bagshaw and Cleland 1990). However, this prevalent overall loosening and tightening will not affect the percentage of standing 5- and 6-day old Arabidopsis hypocotyls in which the zone of gravitropic bending is physically separated from the basal zone that is mostly responsible for the mechanical strength needed for their upright posture (Fig. 2A).

Cellulose is the strongest cell wall component. The requirement of high wall mechanical strength for keeping the shoot upright indicates that the arrangement of microfibrils could affect this process. The modulus of individual cellulose microfibrils ranges from 0.7 to 3.5 GPa (Chanliaud et al. 2002), which is about 100-fold greater than the tensile modulus of primary cell walls (Ryden et al. 2003). Due to this fact, cellulose microfibrils never extend axially under physiological conditions, instead they separate from each other, bend, slide past each other and reorient in the direction of strain during growth, or under external forces, resulting in cell wall deformations (Preston 1982; Refrégier et al. 2004; Zhang et al. 2017). The cellulose mobility depends on microfibril-microfibril and microfibril-matrix interactions, proteinaceous cell wall loosening/tightening factors (Chanliaud et al. 2002; Chanliaud et al. 2004; Zhang et al. 2017) and cellulose orientation that is defined by cortical microtubules (Bringmann et al. 2012; Paredez et al. 2006). BRs affect many processes linked with cellulose. They transcriptionally regulate the majority of *CESA* genes encoding catalytic subunits of cellulose synthase complexes through the involvement of the BR-activated transcription factor BES1 (Xie et al. 2011). Additionally, BRs influence cellulose synthesis post-translationally: the negative regulator of BR signaling, BRASSINOSTEROID INSENSITIVE2 (BIN2), can phosphorylate one of the cellulose synthase subunits that leads to impaired cellulose synthesis (Sánchez-Rodríguez et al. 2017). These mechanisms can be the cause for the BRZ-induced decrease in cellulose content (Fig. 3B). BRs may also change microfibril orientation as they influence cortical microtubule organization (Gupta et al. 2012), possibly via BIN2 (Liu et al. 2018). Hence, the EBL-induced formation of oblique cellulose macrofibrils in the inner wall layer adjacent to the plasma membrane (Fig. 4B, E; Table 1; Supplementary Videos S3, S4) may result from cortical microtubule reorientations.

We observed two additional cellulose-related effects of BRs. First, the BRZ-induced decrease in crystalline cellulose and increase in the TFA-released glucose, without increase in xylose (Fig. 3B, C), can be interpreted as a decrease in the cellulose crystallinity. Similar observations were made in *isoxaben resistant* (*ixr*) mutants (DeBolt et al. 2013). BRZ could thus generate cellulose microfibrils with reduced crystallinity and more exposed glucan chains. This might provide a larger surface area, and thus additional sites, for noncovalent interactions with adjacent microfibrils and matrix polysaccharides. The ‘rough’ surface of such microfibrils could also provide a better access for enzymes forming covalent cross-links between cell wall components. A good candidate for this role is AtXTH3 that catalyses cellulose-cellulose, cellulose-xyloglucan and xyloglucan-xyloglucan transglycosylation (Shinohara et al. 2017). All these additional noncovalent and covalent interactions could explain the increased wall mechanical strength observed with BRZ treatment (Fig. 2B) that contributes to the upright posture of hypocotyls (Fig. 1C).

The second cellulose-related effect of BRs is the EBL-induced macrofibril disorganization at the base of hypocotyls (Fig. 4E; Supplementary Video S4). The overall arrangement of macrofibrils along Arabidopsis hypocotyls is consistent with the classical multinet growth theory, according to which cellulose is deposited transversely next to the plasma membrane, followed by a passive microfibril displacement into deeper wall layers and their axial reorientation, i.e. parallel to the direction of maximal growth (Preston 1982; Refrégier et al. 2004; Roelofsen 1958). As epidermal cells at the base of hypocotyls are at a later developmental stage than those at the apex (Bastien et al. 2016; Gendreau et al. 1997), it means that the majority of longitudinal macrofibrils in the outer wall layer are post-synthetically re-oriented to produce oblique ones in fully elongated cells. A force that drives such re-orientation must be very high. The most likely candidate for this role is a change in the wall hydration generating forces that can rearrange cellulose microfibrils (Huang et al. 2018). In line with this hypothesis, BRs were shown to induce a rapid wall swelling (Caesar et al. 2011). Irrespectively of the mechanisms involved, the elimination and thinning of longitudinal cellulose macrofibrils observed at the base of hypocotyls from EBL-grown plants could decrease their resistance to bending, such that they can easily curve under their own weight and fall on the agar surface (Fig. 1B), which would decrease the percentage of standing hypocotyls. It is, however, important to note that the EBL and BRZ treatment may also affect the content and distribution of other hormones that may contribute to the observed effects.

Mannans are minor hemicelluloses of primary cell walls in angiosperms. Their backbones are either composed of (1→4)-β-D-mannose residues only (pure mannan), or contain (1→4)-β-linked D-glucose and D-mannose residues distributed along the chain in a non-regular fashion (glucomannan). Both polysaccharides can have (1→6)-α-D-galactose substitutions on their backbones, in which case they are referred to as galactomannans and galactoglucomannans, respectively (Schröder et al. 2009). Mannans are considered to play a structural role in cell walls based on experiments with cell wall analogues (Whitney et al. 1998). However, doubts have been cast on the structural functions of mannans in cell walls based on the phenotype of Arabidopsis triple mutant *csla2csla3csla9*. Being deficient in mannan biosynthesis and containing no detectable glucomannans, the mutant plants were phenotypically indistinguishable from their wild type counterparts (Goubet et al. 2009). Our findings that BRZ decreases the content of mannose in cell wall matrix polysaccharides (Fig. 3C), while increasing the gravitropic bending of hypocotyls (Vandenbussche et al. 2011; Fig. 6), and that the triple mutation *csla2csla3csla9* increases the gravitropic curvature (Fig. 6) with no effect on the percentage of upright hypocotyls (Supplementary Fig. S2) show that mannans have a negative effect on gravitropic bending. This role in shoot gravitropism is a new function for mannan polysaccharides. Interestingly, our data are consistent with the fact that gravistimulation of wild type Arabidopsis seedlings was accompanied by a strong downregulation of one gene (*ATCSLA15*) responsible for mannan biosynthesis (Millar and Kiss 2013). Additionally, the level of galactoglucomannans was decreased by about 50% in maize stem pulvini as a result of gravistimulation (Zhang et al. 2011). The negative influence of mannans on shoot gravitropism is interesting from an evolutionary perspective. Aquatic ancestors of terrestrial plants used buoyancy to support their bodies (Hejnowicz 1997) and contained mannans and xylans as principal hemicelluloses (Popper and Tuohy 2010). Interestingly, xyloglucans, a class of hemicelluloses that became prevalent in the primary cell walls of many terrestrial plants, emerged in charophycean green algae (Sørensen et al. 2010), which are considered land plant ancestors (Wickett et al. 2014). It thus appears that xyloglucans are better suited than mannans for land habitats, which could relate to their involvement in plant gravitropism (Velasquez et al. 2019). In this evolutionary context it is important to find out why mannans interfere with gravitropic bending. The precise mechanism of the effect is unknown, but we consider three hypotheses:

1. Glucomannans have less regularly organized backbones than xyloglucans (Schroder et al. 2009) making them insufficiently flexible alone or in a complex with different cell wall components to support the rapid formation of gravitropic bending.
2. Gravitropic bending may involve cellulose reorientations because microtubules, the key determinants of cellulose alignment (Paredez et al. 2006), demonstrated very similar reorientations during gravitropism in different plant organs and species (Blancaflor and Hasenstein 1995; Himmelspach et al. 1999; Zhang et al. 2008). At the same time, mannans were shown to affect cellulose organisation (Yu et al. 2014). Hence, these hemicellulosic polymers may also change cellulose reorientations in the course of gravitropism, which could influence its rate.
3. Changes in mannan synthesis can affect gravitropic bending by modulating the level of reactive oxygen species (ROS). This scenario is possible because GDP-mannose is a common precursor in the biosynthesis of mannans and ascorbate (Sawake et al. 2015). Accordingly, the downregulation of *CSLA* gene expression by gravistimulation (Millar and Kiss 2013) may increase the GDP-mannose pool available for ascorbate biosynthesis. The resulting increase in ascorbate, a strong antioxidant, could modulate ROS levels, which are important for gravitropic responses (Joo et al. 2001; Krieger et al. 2016; Singh et al. 2017; Zhou et al. 2018).

These mechanisms are not mutually exclusive. Studying their contribution to shoot gravitropism will shed light to the role of mannans in cell wall architecture. This could also improve our understanding of key cell wall modifications that allowed successful land colonization by aquatic ancestors of terrestrial plants.

## Materials and Methods

### Plant material and growth conditions

*Arabidopsis thaliana* (L. Heynh.) wild type Columbia-0 and mutant plants were grown on half-strength Murashige and Skoog (MS) medium (pH 5.7) (Duchefa Biochemie) containing 0.68% (w/v) microagar (Duchefa Biochemie). Where indicated, the medium was supplemented with stock solutions of epibrassinolide (0.2 mM in methanol), brassinazole (2 mM in methanol) and oryzalin (500 μM in ethanol), such that their final concentrations were 100 nM, 1 μM and 250 nM, respectively. In each case the medium for control untreated plants was supplemented with corresponding volumes of methanol and/or ethanol.

Surface-sterilized seeds were sown aseptically on large (145×20 mm) round Petri plates (Greiner) containing the above-mentioned media. The seeds were stratified for 2 days at 4°C, and their synchronous germination was induced by exposure to fluorescent white light (150 μmol m^-2^ s^-1^) for 6 h at 21°C. The moment of transfer to light, i.e. the beginning of induction, was taken as zero age for experimental plants. After the 6 h induction period the Petri plates were wrapped in two layers of thick aluminium foil and placed horizontally (or, where indicated, vertically) in an environmentally controlled growth room. Five-day-old etiolated plants grown at 21°C were used in experiments, unless specified otherwise.

### Extensometry

Arabidopsis seedlings for creep tests were placed individually into 1.5 ml Eppendorf test tubes, frozen by plunging the closed tubes into liquid nitrogen, stored at −20°C and used for extensometry within two weeks after freezing. *In vitro* extension of frozen/thawed hypocoyls was measured with a custom-built constant load extensometer (Suslov and Verbelen 2006). A 2- or 5-mm-long segment from the specified region of a hypocotyl was secured between clamps of the extensometer and preincubated in a buffer (20 mM MES-KOH, pH 6.0) in the relaxed state for 2 min. Then, its time-dependent extension (creep) was measured in the same buffer under 625 mg or 750 mg loads for 15 min. The relative creep rate was calculated as described in Vandenbussche et al. (2011).

### Cell wall biochemical analyses

The biochemical analyses of cell wall polysaccharides were performed largely as previously published (Foster et al. 2010). Approximately 300 of 5-day-old seedlings grown on one Petri plate were considered as a biological replicate. Each experiment on the wall biochemistry included three biological replicates, and a total of three independent experiments were performed (i.e. n=9). The seedlings were rapidly harvested to a large volume of 70% EtOH, after which their seed coats were manually removed and discarded. The resulting plant material was transferred to a 2 ml Eppendorf test tube, 1.5 ml of 70% EtOH was added, followed by centrifugation at 10,000 rpm for 5 min discarding the supernatant. One ml of acetone was added to the resulting pellet, followed by centrifugation at 10,000 rpm for 5 min. The supernatant was discarded, and the residue was air-dried overnight. The resulting dry material was ball-milled (Retsch) for 1 min. One ml of EtOH was added to the residue, followed by centrifugation at 21,000 g for 10 min discarding the supernatant. Then 1 ml of MeOH:chloroform (1:1) mixture was added, followed by centrifugation at 21,000 g for 10 min, discarding the supernatant and air-drying the pellet. The resulting powdered cell wall pellet was weighed and transferred to a clean screw-capped 2 ml plastic Sarstedt test tube. *Myo*-Inositol (30 μl of 1 mg ml^-1^ solution in water) was added to the test tube as internal standard. Then 250 μl of 2M TFA was added, followed by incubation at 121°C for 1 h to hydrolyze cell wall polysaccharides. The tube was rapidly cooled on ice, 300 μl of 2-propanol was added and evaporated at 40°C, and the step with 2-propanol was repeated two more times. After this 250 μl of H_2_O was added, and the tube was vortexed, sonicated for 10 min and centrifuged at 21,000 g for 10 min. One part of the resulting supernatant was taken for uronic acids quantification (2 × 50 μl), and the other portion (100 μl) was dried and used for assaying neutral cell wall monosaccharides, while the pellet was processed for crystalline cellulose quantification. Subsequent cell wall biochemical analyses were carried out as described in Sánchez-Rodríguez et al. (2012). Uronic acids were colorimetrically measured using 2-hydroxydiphenyl as reagent (Vilím 1985) with galacturonic acid as standard (Filisetti-Cozzi and Carpita 1991). Cell wall monosaccharides were assayed as alditol acetate derivatives (Neumetzler 2010) using a modified protocol from Albersheim et al. (1967) by gas chromatography performed on an Agilent 6890N GC system coupled with an Agilent 5973N mass selective detector. Crystalline cellulose was determined by Seaman hydrolysis (Selvendran and O’Neill 1987) using Glc equivalents as standard, where the cellulosic Glc content was determined with the anthrone assay (Dische 1962).

### Microscopy

Cellulose macrofibrils were visualized in cell walls with Pontamine Fast Scarlet 4B (S4B) dye (Anderson et al. 2010). In some experiments, living seedlings were stained in 0.2% (w/v) S4B for 30 min, which revealed macrofibrils only in the upper growing parts of hypocotyls. To provide the dye penetration to the cell walls at the base of hypocotyls, plants were extracted under mild conditions by sequential washes with EtOH:100% acetic acid (7:1, v/v) for 1 h; 100% EtOH for 15 min; 50% EtOH in H_2_O for 15 min; and 1M KOH in H_2_O for 30 min. All washing steps were carried out on a rotator. Experimental samples were stored in 1M KOH at 4°C before analysis or used immediately after the last alkaline wash for cellulose visualization. The samples were rinsed in H_2_O before staining to remove residual KOH, after which 0.2% (w/v) S4B was added for 30 min. Then they were rapidly rinsed with a large volume of H_2_O to remove excessive dye and observed under a spinning disc confocal microscope or near-super-resolution Airyscan confocal microscope (Zeiss LSM880). The equipment for spinning disc confocal microscopy included a Nikon Ti-E inverted confocal microscope equipped with a CSU-X1 spinning disc head (Yokogawa, Japan), a 100x CFI Apo oil immersion TIRF objective (NA 1.49, Nikon, Japan), an evolve charge-coupled device camera (Photometrics Technology, USA), and a 1.2x lens between the spinning disc and camera. S4B was excited using a 561 nm laser (similar to Anderson et al. 2010). Image acquisitions were performed using Metamorph software (Molecular Devices, USA) version 7.5. High-resolution imaging of cellulose macrofibrils was performed on a Zeiss LSM880 microscope equipped with an AiryScan Unit (Huff 2015). S4B was excited using a 514 nm Argon Laser through a 458/514 MBS and a 63x Plan-Apochromat oil objective with a numeric aperture of 1.4. The 514 laser was used as 535 nm was indicated as an optimal S4B excitation wavelength (Anderson et al. 2010), which yielded good images for analyses. Emission was detected using the 32 GaAsP PMT array AiryScan unit. For each 8-bit image, a Z-stack consisting of 2-5 images with a 1 μm step-size, an image size of 1672×1672 pixels, and with 1.26 μs pixel dwell time was recorded. The recordings were deconvoluted using the Zeiss ZEN software at highest possible resolution.

Epidermal cell images at the base of hypocotyls were captured with a Leica DM 4000 light microscope equipped with a 1.3 megapixel CCD camera using a dark field mode, under a HCX PL FLUOTAR 20x/0.40 CORR objective. Cell lengths were then measured on digital images using a segmented line tool in ImageJ 1.49b software.

### Time lapse photography by infrared imaging

For gravitropic reorientation assays, plants were grown in infrared light (930 nm) on vertical plates containing the half-strength MS medium with microagar. Plates with three-day-old seedlings were rotated 90° (time point zero). Subsequently, the seedlings were imaged using infrared enabled cameras (Vandenbussche et al. 2010). Images were analysed using the angle tool in ImageJ.

## Acknowledgments

M.S. and S.P. would like to acknowledge the support from the Biological Optical Microscopy Platform (BOMP) at the University of Melbourne. D.S. acknowledges the Research park of Saint Petersburg State University: Center for Molecular and Cell Technologies, and Chromas Core Facility. We are grateful to Professor Paul Dupree (University of Cambridge) for the kind gift of *csla2csla3csla9* seeds.

## References

Albersheim, P., Nevins, D.J., English, P.D. and Karr, A. (1967) A method for the analysis of sugars in plant cell-wall polysaccharides by gas-liquid chromatography. *Acer pseudoplatanus* tissue culture cells. Carbohydr. Res. 5: 340–345.

Anderson, C.T., Carroll, A., Akhmetova, L. and Somerville, C. (2010) Real-time imaging of cellulose reorientation during cell wall expansion in Arabidopsis roots. Plant Physiol. 152: 787–796.

Bagshaw, S.L. and Cleland, R.E. (1990) Wall extensibility and gravitropic curvature of sunflower hypocotyls: correlation between timing of curvature and changes in extensibility. Plant Cell Environ. 13: 85–89.

Bastien, R., Legland, D., Martin, M., Fregosi, L., Peaucelle, A., Douady, S., et al. (2016) KymoRod: a method for automated kinematic analysis of rod-shaped plant organs. Plant J. 88: 468–475.

Blancaflor, E.B. and Hasenstein, K.H. (1995) Time course and auxin sensitivity of cortical microtubule reorientation in maize roots. Protoplasma 185: 72–82.

Bringmann, M., Li, E., Sampathkumar, A., Kocabek, T., Hauser, M.T. and Persson, S. (2012) POM-POM2/cellulose synthase interacting1 is essential for the functional association of cellulose synthase and microtubules in *Arabidopsis*. Plant Cell 24: 163–177.

Caesar, K., Elgass, K., Chen, Z., Huppenberger, P., Witthöft, J., Schleifenbaum, F., et al. (2011) A fast brassinolide-regulated response pathway in the plasma membrane of Arabidopsis thaliana. Plant J. 66: 528–540.

Carpita, N.C. and Gibeaut, D. M. (1993) Structural models of primary cell walls in flowering plants: consistency of molecular structure with the physical properties of the walls during growth. Plant J. 3: 1–30.

Cavalier, D.M., Lerouxel, O., Neumetzler, L., Yamauchi, K., Reinecke, A., Freshour, G., et al. (2008) Disrupting two Arabidopsis thaliana xylosyltransferase genes results in plants deficient in xyloglucan, a major primary cell wall component. Plant Cell 20: 1519–1537.

Chanliaud, E., Burrows, K.M., Jeronimidis, G. and Gidley, M.J. (2002) Mechanical properties of primary plant cell wall analogues. Planta 215: 989–996.

Chanliaud, E., De Silva, J., Strongitharm, B., Jeronimidis, G. and Gidley, M.J. (2004) Mechanical effects of plant cell wall enzymes on cellulose/xyloglucan composites. Plant J. 38: 27–37.

Cosgrove, D.J. (1990a) Rapid, bilateral changes in growth rate and curvature during gravitropism of cucumber hypocotyls: implications for mechanism of growth control. Plant Cell Environ. 13: 227–234.

Cosgrove, D.J. (1990b) Gravitropism of cucumber hypocotyls: biophysical mechanism of altered growth. Plant Cell Environ. 13: 235–241.

Cosgrove, D.J. (2016) Catalysts of plant cell wall loosening [version 1; referees: 2 approved]. F1000Res 5(F1000 Faculty Rev), 119 doi: 10.12688/f1000research.7180.1

Cosgrove, D.J. (2018) Diffuse growth of plant cell walls. Plant Physiol. 176: 16–27.

DeBolt, S., Harris, D. and Stork, J. (2013) Plants and plant products useful for biofuel manufacture and feedstock, and methods of producing same. US 8,383,888 B1.

Dick-Pérez, M., Zhang, Y., Hayes, J., Salazar, A., Zabotina, O.A. and Hong, M. (2011) Structure and interactions of plant cell-wall polysaccharides by two- and three-dimensional magic-angle-spinning solid-state NMR. Biochemistry 50: 989–1000.

Dische, Z. (1962) General color reactions. In Methods in carbohydrate chemistry. Edited by Whistler, R.L. and Wolfrom, M.L. pp. 478–481. Academic Press, New York.

Edelmann, H.G. (1997) Gravistimulated asymmetries in the outer epidermal cell walls of graviresponding coleoptiles. Planta 203(Suppl 1): S123–S129.

Edelmann, H.G. and Samajova, O. (1999) Physiological evidence for the accumulation of restrained wall loosening potential on the growth-inhibited side of graviresponding rye coleoptiles. Plant Biol. 1: 57–60.

Filisetti-Cozzi, T.M. and Carpita, N.C. (1991) Measurement of uronic acids without interference from neutral sugars. Anal. Biochem. 197: 157–162.

Foster, C.E., Martin, T.M. and Pauly, M. (2010) Comprehensive compositional analysis of plant cell walls (lignocellulosic biomass) part II: carbohydrates. J. Vis. Exp. 37: 1837.

Gendreau, E., Traas, J., Desnos, T., Grandjean, O., Caboche, M. and Höfte, H. (1997) Cellular basis of hypocotyl growth in Arabidopsis thaliana. Plant Physiol. 114: 295–305.

Goda, H., Shimada, Y., Asami, T., Fujioka, S. and Yoshida, S. (2002) Microarray analysis of brassinosteroid-regulated genes in Arabidopsis. Plant Physiol. 130: 1319–1334.

Goubet, F., Barton, C.J., Mortimer, J.C., Yu, X., Zhang, Z., Miles, G.P., et al. (2009) Cell wall glucomannan in Arabidopsis is synthesised by CSLA glycosyltransferases, and influences the progression of embryogenesis. Plant J. 60: 527–538.

Gupta, A., Singh, M., Jones, A.M. and Laxmi, A. (2012) Hypocotyl directional growth in Arabidopsis: a complex trait. Plant Physiol. 159: 1463–1476.

Hejnowicz, Z. (1997) Graviresponses in herb and trees: a major role for the redistribution of tissue and growth stresses. Planta 203(Suppl 1): S136–S146.

Himmelspach, R., Wymer, C.L., Lloyd, C.W. and Nick, P. (1999) Gravity-induced reorientation of cortical microtubules observed in vivo. Plant J. 18: 449–453.

Hoson, T. and Wakabayashi, K. (2015) Role of the plant cell wall in gravity resistance. Phytochemistry 112: 84–90.

Huang, S., Makarem, M., Kiemle, S.N., Zheng, Y., He, X., Ye, D., et al. (2018) Dehydration-induced physical strains of cellulose microfibrils in plant cell walls. Carbohyd. Polym. 197: 337–348.

Huff, J. (2015) The Airyscan detector from ZEISS: confocal imaging with improved signal-to-noise ratio and super-resolution. Nat. Methods 12: 1205.

Ikushima, T., Soga, K., Hoson, T. and Shimmen, T. (2008) Role of xyloglucan in gravitropic bending of azuki bean epicotyl. Physiol. Plant. 132: 552–565.

Ivakov, A., Flis, A., Apelt, F., Fünfgeld, M., Scherer, U., Stitt, M., et al. (2017) Cellulose synthesis and cell expansion are regulated by different mechanisms in growing Arabidopsis hypocotyls. Plant Cell 29: 1305–1315.

Joo, J.H., Bae, Y.S. and Lee, J.S. (2001) Role of auxin-induced reactive oxygen species in root gravitropism. Plant Physiol. 126: 1055–1060.

Kiss, J.Z. (2000) Mechanisms of the early phases of plant gravitropism. Crit. Rev. Plant Sci. 19: 551–573.

Krieger, G., Shkolnik, D., Miller, G. and Fromm, H. (2016) Reactive oxygen species tune root tropic responses. Plant Physiol. 172: 1209–1220.

Liu, X., Yang, Q., Wang, Y., Wang, L., Fu, Y. and Wang, X. (2018) Brassinosteroids regulate pavement cell growth by mediating BIN2-induced microtubule stabilization. J. Exp. Bot. 69: 1037–1049.

Millar, K.D. and Kiss, J.Z. (2013) Analyses of tropistic responses using metabolomics. Am. J. Bot. 100: 79–90.

Miller, N.D., Parks, B.M. and Spalding, E.P. (2007) Computer-vision analysis of seedling responses to light and gravity. Plant J. 52: 374–381.

Morita, M.T. (2010) Directional gravity sensing in gravitropism. Annu. Rev. Plant Biol. 61: 705–720.

Neumetzler, L. (2010) Identification and characterization of Arabidopsis mutants associated with xyloglucan metabolism. p. 210. Rhombos-Verlag, Berlin.

Paredez, A.R., Somerville, C.R. and Ehrhardt, D.W. (2006) Visualization of cellulose synthase demonstrates functional association with microtubules. Science 312: 1491–1495.

Park, Y.B. and Cosgrove, D.J. (2012) A revised architecture of primary cell walls based on biomechanical changes induced by substrate-specific endoglucanases. Plant Physiol. 158: 1933–1943.

Pelletier, S., Van Orden, J., Wolf, S., Vissenberg, K., Delacourt, J., Ndong, Y.A., Pelloux, J., et al. (2010) A role for pectin de-methylesterification in a developmentally regulated growth acceleration in dark-grown Arabidopsis hypocotyls. New Phytol. 188: 726–739.

Phyo, P., Wang, T., Xiao, C., Anderson, C.T. and Hong, M. (2017a) Effects of pectin molecular weight changes on the structure, dynamics, and polysaccharide interactions of primary cell walls of Arabidopsis thaliana: insights from solid-state NMR. Biomacromolecules 18: 2937–2950.

Phyo, P., Wang, T., Kiemle, S.N., O’Neill, H., Pingali, S.V., Hong, M., et al. (2017b) Gradients in wall mechanics and polysaccharides along growing inflorescence stems. Plant Physiol. 175: 1593–1607.

Popper, Z.A. and Tuohy, M.G. (2010) Beyond the green: understanding the evolutionary puzzle of plant and algal cell walls. Plant Physiol. 153: 373–383.

Preston, R.D. (1982) The case for multinet growth in growing walls of plant cells. Planta 155: 356–363.

Ray, P.M., Green, P.B. and Cleland, R. (1972) Role of turgor in plant cell growth. Nature 239: 163–164.

Refrégier, G., Pelletier, S., Jaillard, D. and Höfte, H. (2004) Interaction between wall deposition and cell elongation in dark-grown hypocotyl cells in Arabidopsis. Plant Physiol. 135: 959–968.

Roelofsen, P.A. (1958) Cell-wall structure as related to surface growth. Acta Bot. Neer. 7: 77–89.

Ryden, P., Sugimoto-Shirasu, K., Smith, A.C., Findlay, K., Reiter, W.D. and McCann, M.C. (2003) Tensile properties of Arabidopsis cell walls depend on both a xyloglucan cross-linked microfibrillar network and rhamnogalacturonan II-borate complexes. Plant Physiol. 132: 1033–1040.

Sánchez-Rodríguez, C., Bauer, S., Hématy, K., Saxe, F., Ibáñez, A.B., Vodermaier, V., et al. (2012) Chitinase-like1/pom-pom1 and its homolog CTL2 are glucan-interacting proteins important for cellulose biosynthesis in Arabidopsis. Plant Cell 24: 589–607.

Sánchez-Rodríguez, C., Ketelaar, K., Schneider, R., Villalobos, J.A., Somerville, C.R., Persson, S., et al. (2017) BRASSINOSTEROID INSENSITIVE2 negatively regulates cellulose synthesis in *Arabidopsis* by phosphorylating cellulose synthase 1. P. Natl. Acad. Sci. USA 114: 3533–3538.

Sawake, S., Tajima, N., Mortimer, J.C., Lao, J., Ishikawa, T., Yu, X., et al. (2015) KONJAC1 and 2 are key factors for GDP-mannose generation and affect L-ascorbic acid and glucomannan biosynthesis in Arabidopsis. Plant Cell 27: 3397–3409.

Schröder, R., Atkinson, R.G. and Redgwell, R.J. (2009) Re-interpreting the role of endo-beta-mannanases as mannan endotransglycosylase/hydrolases in the plant cell wall. Ann. Bot. 104: 197–204.

Selvendran, R.R. and O’Neill, M.A. (1987) Isolation and analysis of cell walls from plant material. Method. Biochem. Anal. 32: 25–153.

Shah, D.U., Reynolds, T.P.S. and Ramage, M.H. (2017) The strength of plants: theory and experimental methods to measure the mechanical properties of stems. J. Exp. Bot. 68: 4497–4516.

Shinohara, N., Sunagawa, N., Tamura, S., Yokoyama, R., Ueda, M., Igarashi, K., et al. (2017). The plant cell-wall enzyme AtXTH3 catalyses covalent cross-linking between cellulose and cello-oligosaccharide. Sci. Rep. 7: 46099.

Singh, A.P. and Savaldi-Goldstein, S. (2015) Growth control: brassinosteroid activity gets context. J. Exp. Bot. 66: 1123–1132.

Singh, K.L., Mukherjee, A. and Kar, R.K. (2017) Early axis growth during seed germination is gravitropic and mediated by ROS and calcium. J. Plant Physiol. 216: 181–187.

Song, L., Zhou, X.Y., Li, L., Xue, L.J., Yang, X. and Xue, H.W. (2009) Genome-wide analysis revealed the complex regulatory network of brassinosteroid effects in photomorphogenesis. Mol. Plant 2: 755–772.

Sørensen, I., Domozych, D. and Willats, W.G. (2010) How have plant cell walls evolved? Plant Physiol. 153: 366–372.

Sun, Y., Fan, X.Y., Cao, D.M., Tang, W., He, K., Zhu, J.Y., et al. (2010) Integration of brassinosteroid signal transduction with the transcription network for plant growth regulation in Arabidopsis. Dev. Cell 19: 765–777.

Suslov, D. and Verbelen, J.P. (2006) Cellulose orientation determines mechanical anisotropy in onion epidermis cell walls. J. Exp. Bot. 57: 2183–2192.

Suslov, D., Verbelen, J.P. and Vissenberg, K. (2009) Onion epidermis as a new model to study the control of growth anisotropy in higher plants. J. Exp. Bot. 60: 4175–4187.

Tan, L., Eberhard, S., Pattathil, S., Warder, C., Glushka, J., Yuan, C., et al. (2013) An Arabidopsis cell wall proteoglycan consists of pectin and arabinoxylan covalently linked to an arabinogalactan protein. Plant Cell 25: 270–287.

Vandenbussche, F., Petrásek, J., Zádníková, P., Hoyerová, K., Pesek, B., Raz, V., et al. (2010) The auxin influx carriers AUX1 and LAX3 are involved in auxin-ethylene interactions during apical hook development in Arabidopsis thaliana seedlings. Development 137: 597–606.

Vandenbussche, F., Suslov, D., De Grauwe, L., Leroux, O., Vissenberg, K. and Van der Straeten, D. (2011) The role of brassinosteroids in shoot gravitropism. Plant Physiol. 156: 1331–1336.

Velasquez, S.M., Gallemi, M., Aryal, B., Venhuizen, P., Barbez, E., Dünser, K., et al. (2019) Auxin-dependent xyloglucan remodelling defines differential tissue expansion in *Arabidopsis thaliana*. BioRxiv. doi: https://doi.org/10.1101/808964

Vilím, V. (1985) Colorimetric estimation of uronic acids using 2-hydroxydiphenyl as a reagent. Biomed. Biochim. Acta 44: 1717–1720.

Whitney, S.E.C., Brigham, J.E., Darke, A.H., Reid, J.S.G. and Gidley, M.J. (1998) Structural aspects of the interaction of mannan-based polysaccharides with bacterial cellulose. Carbohyd. Res. 307: 299–309.

Wickett, N.J., Mirarab, S., Nguyen, N., Warnow, T., Carpenter, E., Matasci, N., et al. (2014) Phylotranscriptomic analysis of the origin and early diversification of land plants. P. Natl. Acad. Sci. USA 111: E4859–E4868.

Winter, D., Vinegar, B., Nahal, H., Ammar, R., Wilson, G.V. and Provart, NJ. (2007) An “Electronic Fluorescent Pictograph” browser for exploring and analyzing large-scale biological data sets. PLoS One 2: e718.

Xie, L., Yang, C. and Wang, X. (2011) Brassinosteroids can regulate cellulose biosynthesis by controlling the expression of *CESA* genes in *Arabidopsis*. J. Exp. Bot. 62: 4495–4506.

Yu, L., Shi, D., Li, J., Kong, Y., Yu, Y., Chai, G., et al. (2014) CELLULOSE SYNTHASE-LIKE A2, a glucomannan synthase, is involved in maintaining adherent mucilage structure in Arabidopsis seed. Plant Physiol. 164: 1842–1856.

Yu, X., Li, L., Zola, J., Aluru, M., Ye, H., Foudree, A., et al. (2011) A brassinosteroid transcriptional network revealed by genome-wide identification of BESI target genes in Arabidopsis thaliana. Plant J. 65: 634–646.

Zhang, Q., Pettolino, F.A., Dhugga, K.S., Rafalski, J.A., Tingey, S., Taylor, J., et al. (2011) Cell wall modifications in maize pulvini in response to gravitational stress. Plant Physiol. 156: 2155–2171.

Zhang, T., Vavylonis, D., Durachko, D.M. and Cosgrove, D.J. (2017) Nanoscale movements of cellulose microfibrils in primary cell walls. Nat. Plants 3: 17056.

Zhang, Z., Friedman, H., Meir, S., Rosenberger, I., Halevy, A.H. and Philosoph-Hadas, S. (2008) Microtubule reorientation in shoots precedes bending during the gravitropic response of cut snapdragon spikes. J. Plant Physiol. 165: 289–296.

Zhou, L., Hou, H., Yang, T., Lian, Y., Sun, Y., Bian, Z., et al. (2018) Exogenous hydrogen peroxide inhibits primary root gravitropism by regulating auxin distribution during Arabidopsis seed germination. Plant Physiol. Bioch. 128: 126–133.

