## SupplementaryData for "Brassinosteroids Influence Arabidopsis Hypocotyl Graviresponses Through Changes In Mannans And Cellulose": Supplementary Figures and Legends to Supplementary Videos.pdf

**Supplementary Figures S1-S3 and Legends to Supplementary Videos**

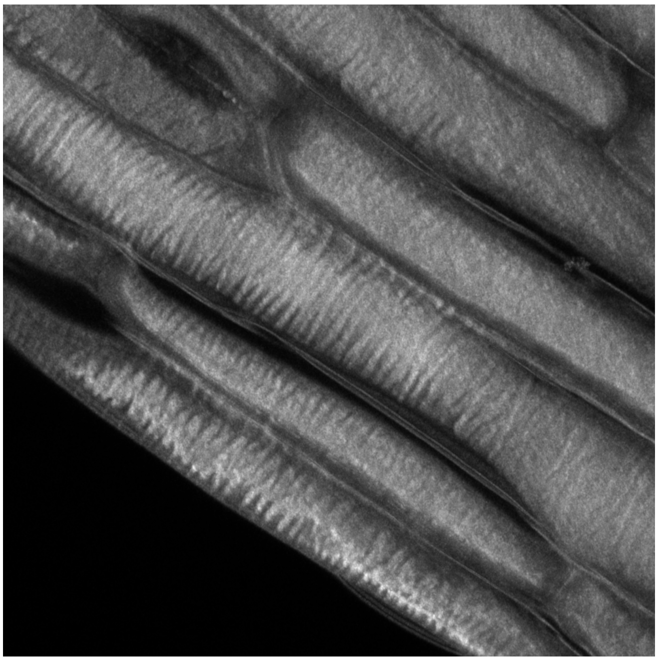

**Fig. S1** Transverse buckling in the innermost layer of outer epidermal cell walls in the upper part of living *Col-0* Arabidopsis hypocotyls. A projection of several optical sections across the whole wall thickness obtained with Airyscan confocal microscope after Pontamine Fast Scarlet 4B staining without preliminary cell wall extraction.

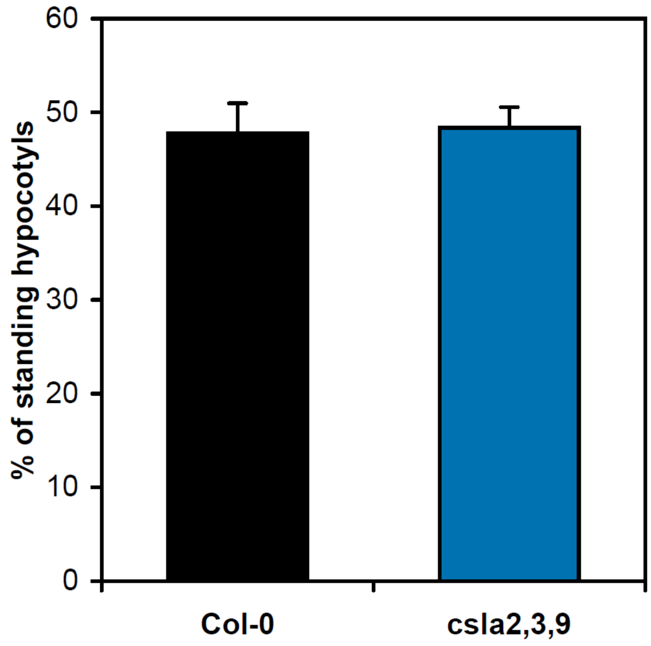


**Fig. S2** The percentage of standing hypocotyls in 5-day-old *Col-0* and *csla2csla3csla9* etiolated Arabidopsis seedlings grown on horizontal Petri plates.

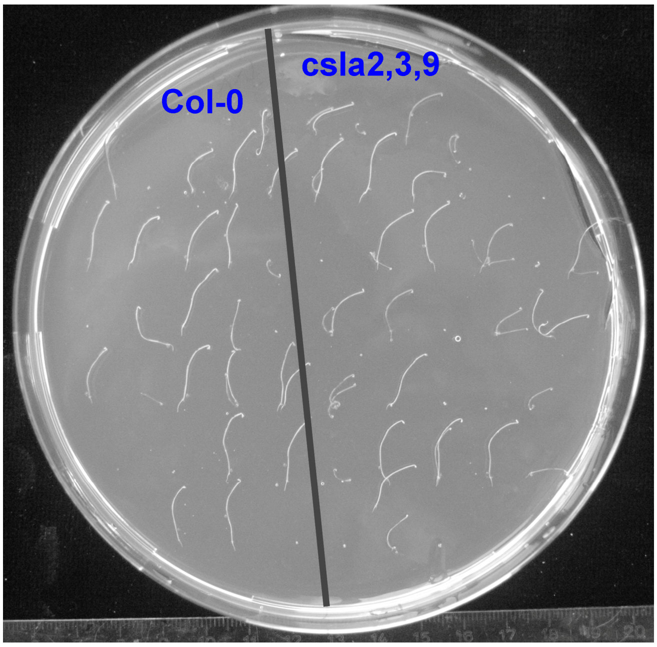


**Fig. S3** Gravitropic bending of hypocotyls in *Col-0* and *csla2csla3csla9* Arabidopsis plants. Etiolated seedlings grown on vertical Petri plates for three days were gravistimulated by a 90-degree counterclockwise rotation of the plates, grown for two more days and photographed. The angles of curvature are 45.0±19.2 degrees (mean ±SD, n=20) for *Col-0* and 64.8±16.5 degrees (mean ±SD, n=21) for *csla2csla3csla9*. This difference is highly significant ($P$ = 0.001; Student's $t$-test).

**Legends to Supplementary Videos**

In all videos cellulose arrangement was observed using spinning disc confocal microscopy. It is represented in the order from the innermost to the outermost layer of the outer epidermal cell wall.

**Video S1** Cellulose macrofibrils in the upper region of hypocotyls from control untreated *Col-0* seedlings.

**Video S2** Cellulose macrofibrils in the basal region of hypocotyls from control untreated *Col-0* seedlings.

**Video S3** Cellulose macrofibrils in the upper region of hypocotyls from EBL-grown *Col-0* seedlings.

**Video S4** Cellulose macrofibrils in the basal region of hypocotyls from EBL-grown *Col-0* seedlings.

**Video S5** Cellulose macrofibrils in the upper region of hypocotyls from oryzalin-grown *Col-0* seedlings.

**Video S6** Cellulose macrofibrils in the basal region of hypocotyls from oryzalin-grown *Col-0* seedlings.

**Video S7** Cellulose macrofibrils in the upper region of hypocotyls from BRZ-grown *Col-0* seedlings.

**Video S8** Cellulose macrofibrils in the upper region of hypocotyls from *Col-0* seedlings grown with BRZ plus oryzalin.

**Video S9** Cellulose macrofibrils in the basal region of hypocotyls from BRZ-grown *Col-0* seedlings.

**Video S10** Cellulose macrofibrils in the basal region of hypocotyls from *Col-0* seedlings grown with BRZ plus oryzalin.
